# Evaluation of the broth microdilution methodology for susceptibility testing of *Mycobacterium tuberculosis* in Peru

**DOI:** 10.1101/2022.03.29.486329

**Authors:** Zully M. Puyen, David Santos-Lázaro, Aiko N. Vigo, Jorge Coronel, Miriam J. Alarcón, Vidia V. Cotrina, David A.J. Moore

## Abstract

Peru is amongst the 30 countries with the highest burden of multidrug-resistant tuberculosis worldwide. In the fight against drug-resistant tuberculosis, the UKMYC6 microtiter plate was developed and validated by the CRyPTIC project. Our objective was to evaluate the use of the broth microdilution UKMYC6 plate for susceptibility testing of drug-resistant *Mycobacterium tuberculosis* (MTB) strains in Peru. 496 nationally-representative MTB strains determined drug-resistant by the routine agar proportion method (APM) were selected. MICs of 13 anti-tuberculosis drugs were determined for each strain using the microdilution UKMYC6 plates and compared with the APM result. MIC distributions for APM-susceptible and APM-resistant strains were demonstrated for rifampicin, isoniazid, kanamycin, and levofloxacin, with reasonable agreement (0.64≤k≤0.79) for rifampicin, ethambutol, ethionamide and kanamycin and the best agreement for isoniazid and levofloxacin (k>0.8). No strain presented MICs higher than the CRyPTIC Epidemiological cut-off values for the new (bedaquiline, delamanid) or repurposed (clofazimine, linezolid) drugs. The microbroth dilution method using the UKMYC6 microtiter plate allowed the complete susceptibility characterization, through the determination of MICs, of drug-resistant *Mycobacterium tuberculosis* strains in Peru. This methodology showed a good diagnostic performance for the drugs rifampicin, isoniazid, kanamycin and levofloxacin drugs.

## INTRODUCTION

Tuberculosis (TB) is a communicable, preventable and curable disease. It is the 13th leading cause of death and the second leading infectious killer after COVID-19 (ranking above HIV/AIDS) (1). It is estimated that in 2019, TB was diagnosed in 10 million (range 8.9-11 million) people and 1.4 million died worldwide from this disease (2). The problem of managing and eliminating TB is further exacerbated by the presence of drug-resistant TB, a major public health problem that threatens progress made in TB care and control worldwide (3). In 2019, about half a million people developed rifampicin-resistant TB, of which 78% were multidrug resistant (MDR-TB) (2). Also, in 2018 it was estimated that 6.2% of MDR-TB cases were extensively drug-resistant (XDR-TB) (3).

Peru has 14% of the estimated cases of tuberculosis in the Region of the Americas, with 27,000 new cases of active disease and 17,000 new cases of smear-positive pulmonary TB each year. In addition, it is one of the 30 countries in the world with the highest burden of MDR-TB (3). TB and MDR-TB is distributed in the 24 departments of Peru; however, the department of Lima (capital of Peru) and the constitutional province of Callao account for 61% of TB cases and 78% of MDR-TB and XDR-TB cases (4).

Different methodologies have been implemented over the years in Peru for the evaluation of resistance to the drugs used in the treatment of tuberculosis. Since 2004 the gold standard method at the National Mycobacterial Reference Laboratory for the *Mycobacterium tuberculosis* complex susceptibility testing has been the 1% indirect APM, which is laborious and requires 2 to 3 weeks from strain inoculation for results (5).

The CRyPTIC (Comprehensive Resistance Prediction for Tuberculosis: An International Consortium) research project has developed and validated the UKMYC6 broth microdilution plate to provide the simultaneous evaluation of MICs of several antituberculosis drugs from a single clinical isolate of MTB. This plate is a variant of the original MYCOTB plate (6, 7), containing 13 different anti-tuberculosis drugs including two repurposed compounds (linezolid and clofazimine) and two new compounds (bedaquiline and delamanid) (8). The original MycoTB microtiter plate showed good results of categorical agreement (92-100%) for the determination of susceptibility to the classic first- and second-line drugs (6, 7) evaluated by the APM method; however, it was only with the development of the UKMYC5 plate that the incorporation of new and redefined drugs was achieved (8). Finally, after the evaluation of this plate, the drug para-aminosalicylic acid was eliminated, optimizing the rest of the concentrations, giving rise to the UKMYC6 plate. In this way, the UKMYC6 microtiter plates provide quantitative values of MICs giving a better understanding of drug resistance in order to provide adequate and individualized treatment to each patient.

The objective of this study was to take advantage of the opportunity presented by the CRyPTIC study of genomic determinants of drug resistance to evaluate the performance of the broth microdilution methodology using UKMYC6 plate for susceptibility testing of MTB strains to anti-tuberculosis drugs in Peru when compared with the APM results. Furthermore, by selection of a nationally representative sample of strains, the data will contribute an overview of the profile of TB drug MICs in drug resistant strains nationally.

## METHODS

### Design, settings and selection of MTB strains

The study was carried out by the National Reference Laboratory for Mycobacteria (Laboratorio de Referencia Nacional de Micobacterias - LRNM) of the National Institute of Health (Instituto Nacional de Salud - INS) in collaboration with investigators at Universidad Peruana Cayetano Heredia (UPCH) and the London School of Hygiene and Tropical Medicine (LSHTM). 496 drug resistant MTB strains (according to routine APM results), representative from all over Peru were selected. These strains had been previously isolated from samples of patients with pulmonary and/or extrapulmonary TB during the years 2015-2018 and were stored in the LRNM culture bank. Each of the selected strains had preliminary information on resistance profiles obtained by the 7H10 APM, and were randomly selected by stratified sampling according to the prevalence of MDR-TB in each one of the 24 departments (in addition to the constitutional province of Callao) of Peru, reported in the mentioned period (4). The results of the routine drug susceptibility tests were obtained from the database exported from the national information system, NETLab, of the INS.

### Ethics statement

Approval for the use and processing of the strains was obtained from the INS Institutional Research Ethics Committee (Reference number OC-020-19). The CRyPTIC associated study (9) – for which the objective was identification of the genomic determinants of MTB drug resistance by Whole Genome Sequencing (WGS) – was reviewed and approved by the Research Ethics Committees of the INS, UPCH and LSHTM. All personal information was anonymized for this analysis.

### Routine susceptibility testing (APM testing)

The susceptibility tests were carried out under programmatic conditions by the LRNM in the period 2015-2018 using the APM on 7H10 agar. The procedures established by the Clinical and Laboratory Standards Institute (CLSI) (10) were followed and the phenotypic susceptibility was determined for the drugs rifampicin, isoniazid, ethambutol, ethionamide, kanamycin and levofloxacin, according to the critical concentrations (CC) recommended by the WHO at the date of testing (Table 1). Briefly, four quadrant Petri dishes containing Middlebrook 7H10 medium (Becton-Dickinson, Sparks, Md., USA) were used. MTB cultures on Lowenstein Jensen were transported to the LRNM from regional laboratories. The strains were subcultured in Middlebrook 7H9 broth (Becton Dickinson, Sparks, USA) and incubated for 7 days at 37°C. Subsequently, fresh broth cultures were standardized to a McFarland 0.5 turbidity scale. The standardized suspensions were diluted 10^-2^ to allow the growth of countable colonies for interpretation. For each culture, the drug-containing quadrants, as well as the drug-free control quadrant, were inoculated with 100 μL of the diluted suspension. The plates were sealed in plastic bags and incubated at 37°C for 21 days. The cultures were classified as resistant when the number of colonies developed in the drug quadrant was more than 1% of the number of colonies observed in the control quadrant, otherwise they were classified as susceptible.

**Table 1:**
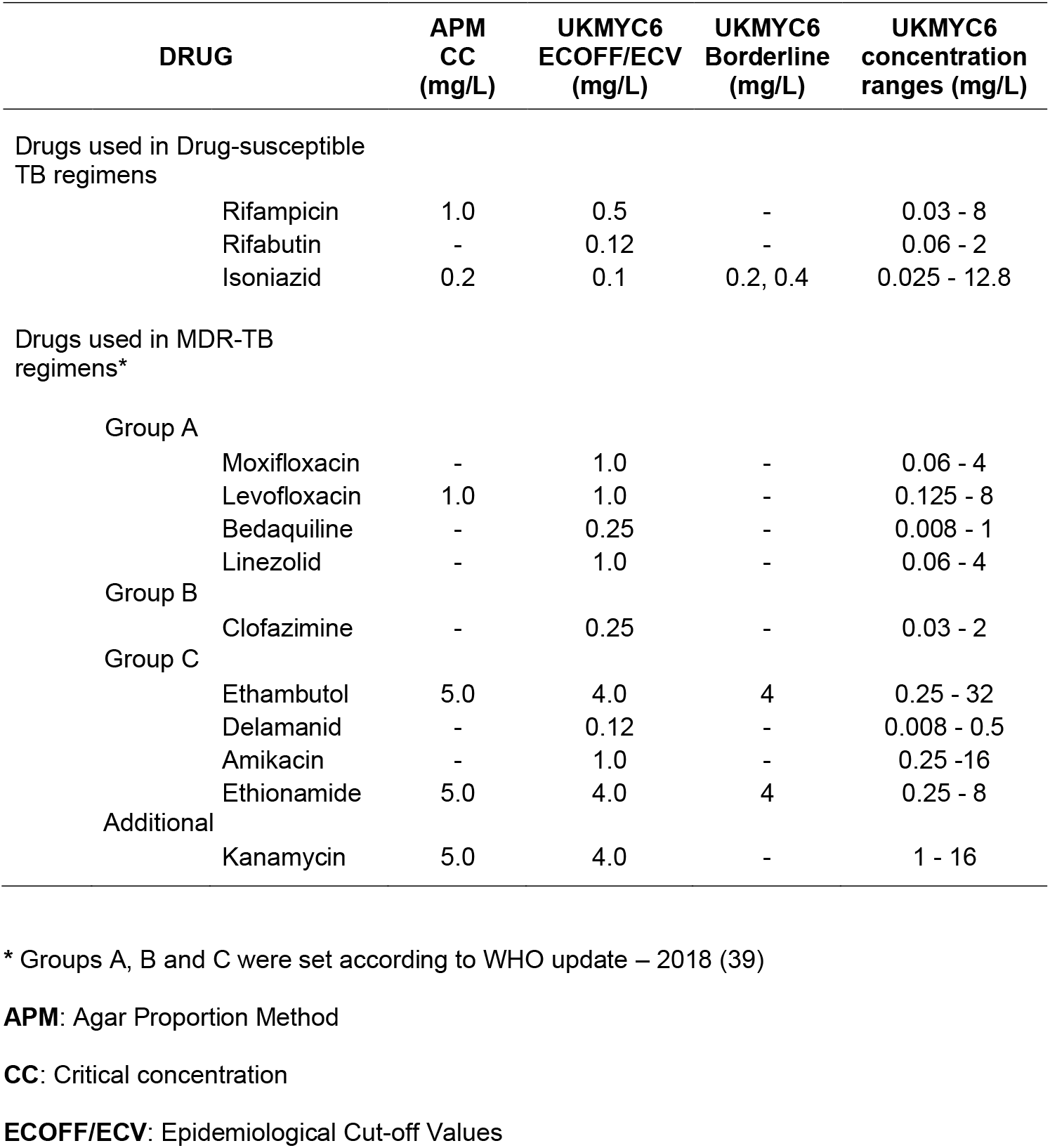
Critical concentrations of APM and the UKMYC6 plate.

| DRUG | APM<br>CC<br>(mg/L) | UKMYC6<br>ECOFF/ECV<br>(mg/L) | UKMYC6<br>Borderline<br>(mg/L) | UKMYC6<br>concentration<br>ranges (mg/L) |
| --- | --- | --- | --- | --- |
| Drugs used in Drug-susceptible TB regimens |  |  |  |  |
| Rifampicin | 1.0 | 0.5 | - | 0.03 - 8 |
| Rifabutin | - | 0.12 | - | 0.06 - 2 |
| Isoniazid | 0.2 | 0.1 | 0.2, 0.4 | 0.025 - 12.8 |
| Drugs used in MDR-TB regimens* |  |  |  |  |
| Group A |  |  |  |  |
| Moxifloxacin | - | 1.0 | - | 0.06 - 4 |
| Levofloxacin | 1.0 | 1.0 | - | 0.125 - 8 |
| Bedaquiline | - | 0.25 | - | 0.008 - 1 |
| Linezolid | - | 1.0 | - | 0.06 - 4 |
| Group B |  |  |  |  |
| Clofazimine | - | 0.25 | - | 0.03 - 2 |
| Group C |  |  |  |  |
| Ethambutol | 5.0 | 4.0 | 4 | 0.25 - 32 |
| Delamanid | - | 0.12 | - | 0.008 - 0.5 |
| Amikacin | - | 1.0 | - | 0.25 - 16 |
| Ethionamide | 5.0 | 4.0 | 4 | 0.25 - 8 |
| Additional |  |  |  |  |
| Kanamycin | 5.0 | 4.0 | - | 1 - 16 |

\* Groups A, B and C were set according to WHO update – 2018 (39)
**APM:** Agar Proportion Method
**CC:** Critical concentration
**ECOFF/ECV:** Epidemiological Cut-off Values

### Reculture and susceptibility testing (microtitre plate testing)

Drug susceptibility testing was performed using the UKMYC6 microtitre panel, designed by the CRyPTIC project (Thermo Fisher Inc., UK). The UKMYC6 plate allowed for the determination of susceptibility against 13 antituberculosis drugs composed of agents used in drug-susceptible TB treatment (rifampicin, rifabutin, isoniazid) as well as longer MDR-TB treatment corresponding to groups A (levofloxacin, moxifloxacin, bedaquiline, linezolid), B (clofazimine), C (ethambutol, delamanid, amikacin, ethionamide) and kanamycin (Table 1). Each drug had 5 to 10 concentrations obtained by serial doubling dilution (Figure S1). The strains that were cryopreserved were reactivated in 7H9 liquid medium, supplemented with oleic acid albumin dextrose catalase (OADC) (Thermo Fisher, Scientific Inc., USA), for 7 days at 37°C. These were then subcultured in Middlebrook 7H10 media for 25 to 30 days at 37°C. From the solid cultures in 7H10 medium, 0.5 McFarland scale suspensions were prepared in Tween saline with glass beads (Thermo Fisher, Scientific Inc., USA). Then, 100μL of the suspension was diluted in a 7H9 broth tube supplemented with OADC to give an inoculum of 1×10^5^ CFU/mL (range: 5×10^4^ to 5×10^5^ CFU/mL). Subsequently, using the automated Sensititre Autoinoculator®/AIM^®^ equipment (Thermo Fisher, Scientific Inc., USA), 100μL of inoculum was dispensed into each well of the UKMYC6 plate. The plates were sealed using clear plastic and incubated aerobically at 35-37°C. The H37Rv ATCC 27294 strain was used to perform periodic quality control tests of the analyzed drugs, as well as to quality control for contamination and adequate growth in two positive control (drug-free) wells of each plate used. All laboratory work related to the culture of live bacteria was carried out in the biosafety level 3 facilities of the INS and the UPCH.

### Determination of MICs

Plates were read using the semi-automated Vizion™ instrument. The results of the plates were considered valid only when the positive control wells showed acceptable growth and free of contamination. The plates were read 14 days after inoculation, as established by the CRyPTIC project (8). Additionally, if the growth of the positive control was weak or insufficient, a second reading was carried out at 21 days. The Vizion™ system captured and stored an image of the recorded MICs of each plate. The MIC of a drug was considered as the lowest concentration capable of inhibiting the visible growth of a microorganism in a given well.

### DNA extraction and genomic sequencing

WGS was performed for all strains for the primary objective of the parent trial. For this study, genomic results were only used for discrepant analysis to resolve discordant results between the APM and microtitre tests. DNA extraction and microtitre susceptibility testing were performed starting from the same solid culture (7H10 culture) to minimize phenotypic and genotypic variation. Genomic DNA was isolated using the phenol-chloroform method (Supplementary method). Sequencing libraries were generated using the Nextera XT Library Preparation Kit, following the manufacturer’s recommendations. WGS was performed at the University of Oxford (UK) using the Illumina HiSeq platform (Illumina Inc., San Diego, CA, USA). Paired end 150 bp sequencing reads were generated and stored in fastq.gz files.

### Bioinformatic Analysis

The quality of the sequencing reads was evaluated using the FastQC v0.11.9 (11) program. The paired reads were filtered with the Trimmomatic v0.38 (12) program using default values and a minimum Phred score of 20. The filtered reads were mapped against the H37Rv reference genome (NC_000962.3) using BWA v0.7.17 (13). The elimination of duplicate readings was carried out with the program Picardtools v2.18.25 (14). The variant call was made using the GATK v4 (15) program. Resistance-associated genes were evaluated for genotypic resistance to rifampin (*rpoB*), isoniazid (*inhA, katG, ahpC, fabG1, kasA*), fluoroquinolones (*gyrA, gyrB*), and second-line injectables (*rrs*, *eis, tlyA*). The genetic variants found were validated using the list of mutations published by the WHO in the “Catalogue of mutations in *Mycobacterium tuberculosis* complex and their association with drug resistance” (16). Likewise, the resistance profiles were determined using the programs TBProfiler v3.0.4 (17) (database of mutations v. A2a234b) and Mykrobe v0.10 (18). For each drug, the strains were classified as susceptible or resistant according to the absence or presence of mutations detected in the genes associated with resistance, respectively.

### Statistical analysis

Descriptive analyses of the MICs obtained and development times were performed on the UKMYC6 plates. Based on the MICs of the UKMYC6 plate, strains were categorized as susceptible, intermediate or resistant (ternary categorization) taking as reference the Epidemiologic Cut-off Values (ECOFF/ECV), as well as borderline concentrations (concentrations at which genetic mutations present different effects giving rise to mixed categories of susceptibility or resistance), established by the CRyPTIC project (19). A result was determined as susceptible if the MIC was less than or equal to the established ECOFF/ECV; otherwise, it was defined as resistant. Isoniazid, ethambutol and ethionamide were categorized taking into account borderline concentrations (Table S1). The sensitivity, specificity, positive predictive value, negative predictive value and categorical agreement were determined taking as reference values the results of the APM method and comparing them to the susceptible/resistant categorization from the UKMYC6 plate. The calculations of diagnostic performance and determination of Cohen’s Kappa coefficient (k) were performed in the program R v4.0.5 (20) using the packages epiR v2.0.19 (https://cran.r-project.org/web/packages/epiR) and vcd v1.4 (https://cran.r-project.org/web/packages/vcd), respectively. The strength of agreement was established as insignificant (0≤k≤0.2), medium (0.2<k≤0.4), moderate (0.4<k≤0.6), good (0.6<k≤0.8) or almost perfect (0.8<k≤ 1) according to the previously proposed classification (21).

### Data availability

The raw WGS data used in discrepant analysis will shortly be submitted to one of the three main databases.

## RESULTS

In the present study, 496 MTB drug-resistant strains were analysed, of which 70% (347/496) were from Lima and Callao, while the rest came from the remaining 23 departments of Peru in the range of 1 to 18 strains per department (Table S2).

### Phenotypic resistance by APM

The percentages of strains with APM-defined phenotypic resistance included in the study for each one of the drugs were: 86% (427/496) resistant to rifampicin, 94% (464/496) to isoniazid, 45% (218/489) to ethambutol, 31% (155/493) to ethionamide, 11% (56/495) to kanamycin and 11% (54/486) to levofloxacin. In addition, according to the classification of drug resistance profile of the year 2020 (2), 5% (24/496) presented rifampicin mono-resistant TB (RR-TB), 12% (61/496) isoniazid mono-resistant (HR-TB), 76% (376/496) MDR-TB, with a further 5% (27/496) XDR-TB, and 2% (8/496) had different patterns of drug resistance.

### MIC determination using UKMYC6 plate

The microbiological evaluation by the microtitre system using the UKMYC6 plate determined that 80% (397/496) presented final growth readings at 14 days and 20% (99/496) at 21 days. Overall, 99% of the readings obtained for each of the drugs showed valid development results (MICs inside and outside evaluated ranges). Of these, an average of 12% of MIC readings were above the dilution range of the UKMYC6 plate, most frequently rifampicin (68%, 326/480) and rifabutin (46%, 229/494) (Table S3).

### Comparison between APM and microtiter method

Comparative analysis was performed between the APM phenotypic test results and UKMYC6 MICs for 6 drugs: rifampicin, isoniazid, ethambutol, ethionamide, kanamycin and levofloxacin. The phenotypically susceptible or resistant strains by APM were graphed separately by means of histograms showing the MICs values obtained from the UKMYC6 plates (Figure 1A). The drugs rifampicin, isoniazid, kanamycin, and levofloxacin showed visibly different MIC distributions for APM-susceptible and resistant strains. However, there was considerable overlap of the UKMYC6 MIC distribution for strains defined as resistant and susceptible to ethambutol and ethionamide by APM (Figures 1A and S2). For the remaining drugs (rifabutin, amikacin, moxifloxacin, bedaquiline, delamanid, clofazimine, and linezolid) no reference drug susceptibility test (susceptible/resistant) assignment result was available so the MIC distributions for all strains were plotted together. The drugs rifabutin, amikacin and moxifloxacin showed the presence of strains with high and low MICs values for which the use of ECOFF/ECVs established the presence of 68% (336/494), 6% (29/491) and 9% (43/495) prevalence of resistant strains, respectively. However, none of the strains presented MICs higher than the ECOFF/ECVs values for the new or repurposed drugs (bedaquiline, delamanid, linezolid, clofazimine), so they were defined as phenotypically susceptible strains by the UKMYC6 microtitre method (Figure 1B).

**Figure 1:**
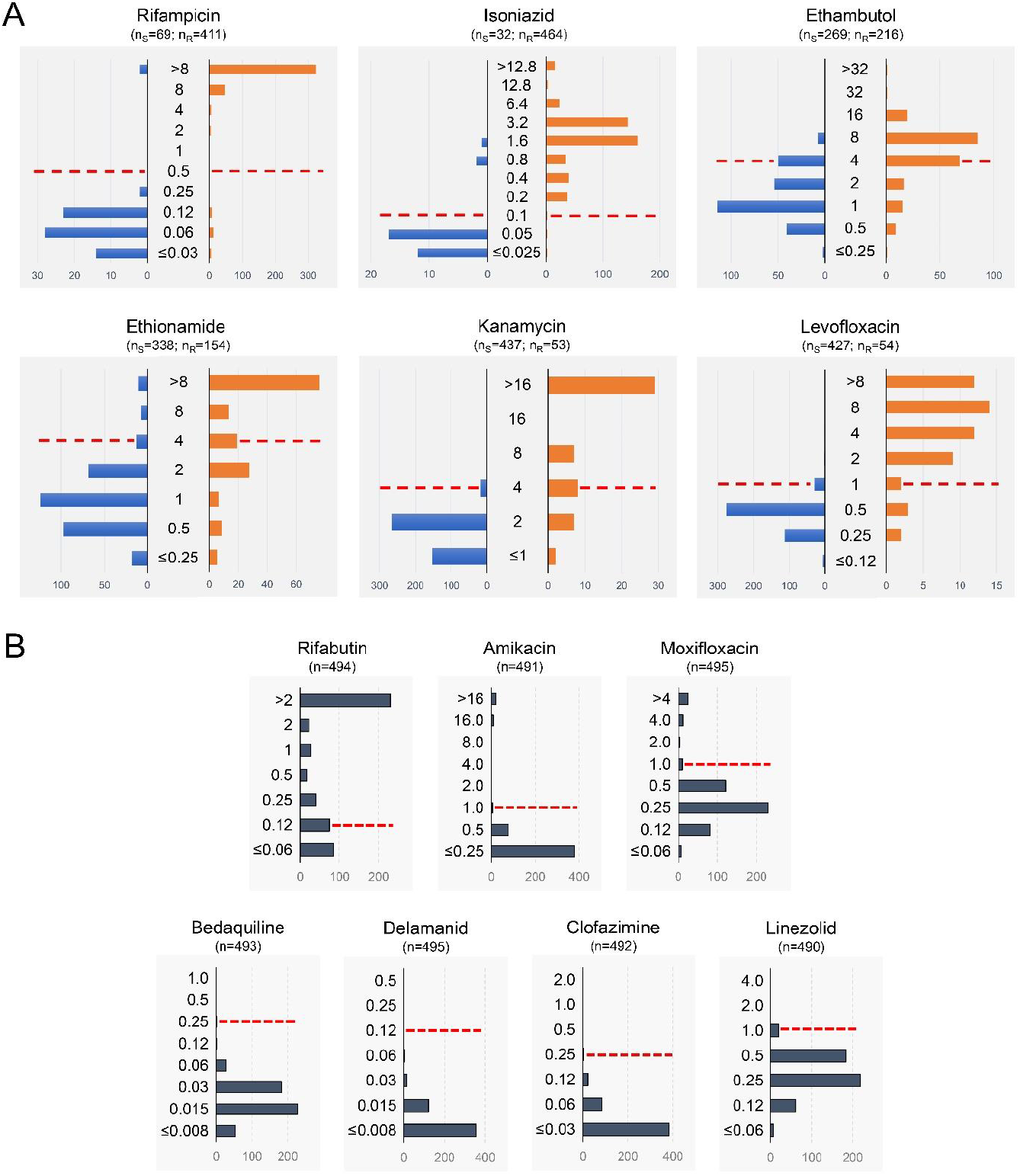
Distribution of MICs in comparison to the results obtained by APM. A) Analysis obtained from the comparison of susceptibility results by APM, categorized as susceptible (blue bars) or resistant (orange bars), in comparison with the MICs obtained in UKMYC6 plates. B) Distribution of MICs of drugs analysed only by the microtiter method. For both analyses, the ECOFF/ECVs of the UKMYC6 plates are indicated by dashed lines. The amounts of susceptible (nS) and resistant (nR) strains per APM are specified for each drug. For the case of drugs that were not evaluated by APM, the total number of strains or measurements (n) performed was specified.

The isoniazid, ethambutol, and ethionamide drugs presented results with borderline MICs in 16% (77/496), 25% (121/492), and 6% (32/495) of their results, respectively. In the case of isoniazid, all borderline MICs were determined in strains categorized as resistant by APM; while in the cases of ethambutol and ethionamide, borderline MICs were determined in both susceptible and resistant strains (Table 2).

**Table 2:** Phenotypic characterization by APM of results with borderline MICs values.

| Drug | APM<br>CC<br>(mg/L) | Borderline<br>MIC<br>(mg/L) | S | APM<br>R | NA | Total |
| --- | --- | --- | --- | --- | --- | --- |
| Isoniazid | 0.2 | 0.2 | 0 | 37 | 0 | 77 |
|  |  | 0.4 | 0 | 40 | 0 |  |
| Ethambutol | 5 | 4 | 50 | 68 | 3 | 121 |
| Ethionamide | 5 | 4 | 13 | 19 | 0 | 32 |

**APM:** Agar Proportion Method.
**CC:** Critical concentration.
**MIC:** Minimum Inhibitory Concentration.
**R:** Number of phenotypically resistant strains.
**S:** Number of phenotypically susceptible strains.
**NA:** Not available.

### Performance of the UKMYC6 microtiter plate compared to APM for categorical susceptible/resistant determination

Using the UKMYC6 ECOFF/ECVs values to categorise strains as susceptible or resistant and comparing these data with APM results for the six drugs with drug susceptibility results in both systems, resulted in an average categorical agreement of 0.93. The highest categorical concordances were obtained for rifampicin, isoniazid, kanamycin and levofloxacin (Table 3). Using APM category as the reference, the sensitivity of UKMYC6 for detection of drug resistance was highest for isoniazid (0.98) and rifampicin (0.93) and lowest for ethionamide (0.66) and kanamycin (0.68). Specificity was high for all drugs, from 0.91 for isoniazid to 1.0 for kanamycin.

**Table 3:** Performance of microtiter method (UKMYC6 MICs) compared to APM for detecting drug resistance.

| Drug | UKMYC6 | APM |  | Sensitivity<br>(95% CI) | Specificity<br>(95% CI) | PPV<br>(95% CI) | NPV<br>(95% CI) | Categorical<br>agreement | Cohen's<br>kappa |
| --- | --- | --- | --- | --- | --- | --- | --- | --- | --- |
|  |  | R | S |  |  |  |  |  |  |
| Rifampicin | R | 383 | 2 | 0.93 (0.90, 0.95) | 0.97 (0.90, 1.00) | 0.99 (0.98, 1.00) | 0.71 (0.60, 0.79) | 0.94 | 0.78 |
|  | S | 28 | 67 |  |  |  |  |  |  |
| Isoniazid | R | 381 | 3 | 0.98 (0.97, 0.99) | 0.91 (0.75, 0.98) | 0.99 (0.98, 1.00) | 0.83 (0.66, 0.93) | 0.98 | 0.85 |
|  | S | 6 | 29 |  |  |  |  |  |  |
| Ethambutol | R | 107 | 8 | 0.72 (0.64, 0.79) | 0.96 (0.93, 0.98) | 0.93 (0.87, 0.97) | 0.84 (0.79, 0.88) | 0.87 | 0.71 |
|  | S | 41 | 211 |  |  |  |  |  |  |
| Ethionamide | R | 89 | 18 | 0.66 (0.57, 0.74) | 0.94 (0.91, 0.97) | 0.83 (0.75, 0.90) | 0.87 (0.83, 0.90) | 0.86 | 0.64 |
|  | S | 46 | 307 |  |  |  |  |  |  |
| Kanamycin | R | 36 | 0 | 0.68 (0.54, 0.80) | 1.00 (0.99, 1.00) | 1.00 (0.90, 1.00) | 0.96 (0.94, 0.98) | 0.97 | 0.79 |
|  | S | 17 | 437 |  |  |  |  |  |  |
| Levofloxacin | R | 47 | 3 | 0.87 (0.75, 0.95) | 0.99 (0.98, 1.00) | 0.94 (0.83, 0.99) | 0.98 (0.97, 0.99) | 0.98 | 0.89 |
|  | S | 7 | 424 |  |  |  |  |  |  |

**APM:** Agar Proportion Method
**R:** Number of phenotypically resistant strains.
**S:** Number of phenotypically susceptible strains.
**PPV:** positive predictive value.
**NPV:** negative predictive value.
**95% CI:** 95% confidence Interval.

### WGS used for the analysis of discordant results

179 discordant results were identified for all six drugs tested by UKMYC6 and APM. There were no ‘UKMYC6-resistant/APM-susceptible’ discrepancies for kanamycin but for all other agents there were discordant results in both directions. Results with intermediate MICs were not considered. Ethambutol and ethionamide had the highest degrees of disagreement with 13% (49/371) and 14% (64/463), respectively. The results of genotypic susceptibility testing by WGS gave similar levels support to the results obtained by APM (52%, 93/179) and those obtained by the UKMYC6 system (48%, 86/179) (Table 4).

**Table 4:** Analysis of discordant results between APM and UKMYC6 methodologies. The genotypic result obtained by WGS is shown. Results with borderline MICs are ignored.

| Drug | Total | APM | UKMYC6 | WGS |  | APM results supported by WGS (%) | UKMYC6 results supported by WGS (%) |
| --- | --- | --- | --- | --- | --- | --- | --- |
|  |  |  |  | S | R |  |  |
| Rifampicin | 30 | S | R | 1 | 1 | 22 (73) | 8 (27) |
|  |  | R | S | 7 | 21 |  |  |
| Isoniazid | 9 | S | R | 2 | 1 | 4 (44) | 5 (56) |
|  |  | R | S | 4 | 2 |  |  |
| Ethambutol | 49 | S | R | 0 | 8 | 23 (47) | 26 (53) |
|  |  | R | S | 18 | 23 |  |  |
| Ethionamide | 64 | S | R | 7 | 11 | 29 (45) | 35 (55) |
|  |  | R | S | 24 | 22 |  |  |
| Kanamycin | 17 | S | R | 0 | 0 | 9 (53) | 8 (47) |
|  |  | R | S | 8 | 9 |  |  |
| Levofloxacin | 10 | S | R | 1 | 2 | 6 (60) | 4 (40) |
|  |  | R | S | 2 | 5 |  |  |

**APM:** Agar Proportion Method.
**WGS:** Whole Genome Sequencing.
**R:** Number of resistant strains.
**S:** Number of susceptible strains.

The use of the borderline MIC criteria for ethambutol considerably decreased the number of discordant results, specifically ‘UKMYC6-susceptible/APM-resistant’ strains, excluding 68 discordant results. However, all the excluded results presented high-reliability mutations associated with resistance. In contrast, for ethionamide 19 ‘UKMYC6-susceptible/APM-resistant’ discordant results were excluded, of which 47% (9/19) presented mutations associated with resistance (Table 4, Table S4).

## DISCUSSION

This is the first description of the comprehensive drug resistance profile, defined by MIC distribution, of a nationally representative sample of strains of *M. tuberculosis* in Peru. Such analyses are of fundamental importance when considering the local design of standardized treatment regimens for MDR-TB (22).

Overall, there was almost no resistance to the new and repurposed drugs identified with no strains exceeding the ECOFF/ECV for bedaquiline, delamanid, clofazimine or linezolid. This results agree with previous studies in the Americas region (23) as well as in other contexts (24, 25). This reflects the sparse usage of these agents within a compassionate use framework prior to their incorporation into national guidelines in 2018 (26) and indicates their introduction into a favourable environment from that timepoint onwards; comparison now with a similarly representative sample of contemporary MDR strains would be instructive and important.

The Sensititre microtitre plate is a convenient tool for the analysis of MICs to any of the drugs used in the treatment of TB (6, 27, 28). The methodology facilitates addition of new drugs and the range of MICs being tested can be adapted, if necessary, in certain settings (8). The WHO-recommended critical concentrations for new and reproposed drugs are still provisional (29, 30) and further work to define MIC distributions in a diversity of settings can contribute to refinement (31, 32). In the Sensititre UKMYC6 plates used (6) for this analysis we found that, for rifampicin-resistant strains, most MICs were out of range for the first- and second-line drugs that have been used in Peru, whilst in contrast no strains exhibited MICs above the ECOFF for the new agents.

The study provided the first opportunity to compare indirect drug susceptibility test (DST) by the proportion method on agar with MIC testing using liquid culture in the Sensititre platform in representative sample of drug-resistant strains of *M. tuberculosis* in Perú. Likewise, once again demonstrating the considerable challenge of defining a binary susceptible/resistant phenotype for certain drugs, most notably, and not unexpectedly, ethambutol and ethionamide (33–35). For levofloxacin and kanamycin, discrepant analysis with WGS found in favour of the proportion method and Sensititre with almost equal frequency. These findings highlight the well-recognised imperfections of all approaches to *M. tuberculosis* DST. Sometimes there is no ‘one right answer’.

What is the clinician to make of the information provided by the laboratory and how should the laboratory present it? There is a reluctance to share MIC data with clinicians who lack the training to interpret it. Few laboratory scientists and even fewer clinicians understand the complexities of the pharmacokinetics of TB drugs and how this relates to the MIC for a particular drug for a particular strain, so it still seems reasonable to try to simplify the message to the binary susceptible/resistant call where possible.

The value of MIC data lies in understanding the drift in the distribution in well characterised populations over time (public health usage) and in case management when therapeutic options are very limited but dosage increases might facilitate efficacy (clinical usage) (36).

An important strength of this analysis is the national representativity. All strains identified nationally during the study period should have been sent to the National Mycobacteria Reference Laboratory for further testing so stratified sampling of the strain bank according to MDR-TB burden during the study period ensured a comprehensive and proportionate national coverage. The availability and use of WGS for discrepant analysis was a critically important enhancement that reinforced the importance of not depending upon a single methodology as the ‘gold standard’. A limitation of the analysis was the lack of proportion method data for the new and repurposed agents, reflecting the earlier time period during which the original proportion method testing was done, and highlighting the power and versatility of a microtitre-based assay in accommodating a large number of drugs within a single assay.

The rapid expansion of the use of WGS for TB DST (37, 38), the growing library of identified resistance-conferring SNPs for all drugs and the tumbling cost (sequencing of the MTB genome is now no more expensive than MGIT phenotypic DST for 4 agents in the UK), place WGS as a likely near-horizon successor to phenotypic DST in settings where the infrastructure allows. Web-based tools for WGS interpretation can deliver (almost instantaneously) a ‘resistance probability’ for *every* drug based upon SNP identification in an uploaded sequence. This is derived by comparison with a large iterative database of paired phenotypic-genotypic data combined with some prediction modelling. Crucially for the clinician, the ‘probability’ acknowledges the uncertainty inherent in the result, allowing for a more intelligent and informed decision-making.

In conclusion, the susceptibility determination system by the microtitre method using the UKMYC6 plate allowed the complete susceptibility characterization, through the determination of MICs, of drug-resistant *Mycobacterium tuberculosis* strains in Peru. Additionally, the analysis of new and redefined drugs was included. Finally, this system showed a good diagnostic performance for the drugs rifampicin, isoniazid, kanamycin and levofloxacin, in comparison with the reference method.

## ACKNOWLEDGEMENTS

We express our thanks to all staff of the NRLM and to the Peruvian network of tuberculosis laboratories, for the routine work in the isolation and identification of different strains that were included in this study.

This work was supported by the Peruvian National Institute of Health and PROCIENCIA (contract N° 230-2018-FONDECYT); the ‘Dirección de Investigación de la Universidad Peruana de Ciencias Aplicadas’, Lima-Peru (A-055-2021-2); Newton Fund Institutional Links award (414591184 Moore PER); Wellcome Trust/Newton Fund-MRC Collaborative Award (200205/Z/15/Z to CRyPTIC) and Bill and Melinda Gates Foundation (OPP1133541 to CRyPTIC).

Conceptualization: Z.M.P., D.A.J.M., D.S.L., J.C.; Project administration: Z.M.P. and D.A.J.M. Methodology: Z.M.P., D.S.L., A.N.V., J.C., M.J.A., V.V.C., D.A.J.M. Data analysis and writing of the manuscript: Z.M.P., D.A.J.M., D.S.L. and J.C. All authors read and approved the final manuscript. The authors declare no conflict of interest.

